# Small molecule inhibition of PIKFYVE kinase rescues gain- and loss-of-function *C9ORF72* ALS/FTD disease processes *in vivo*

**DOI:** 10.1101/685800

**Authors:** K. A. Staats, C. Seah, A. Sahimi, Y. Wang, N. Koutsodendris, S. Lin, D. Kim, W-H. Chang, K. A. Gray, Y. Shi, Y. Li, M. Chateau, V. R. Vangoor, K. Senthilkumar, R. J. Pasterkamp, P. Cannon, B.V. Zlokovic, J. K. Ichida

**Affiliations:** Department of Stem Cell Biology and Regenerative Medicine, Eli and Edythe Broad Center for Regenerative Medicine and Stem Cell Biology, Keck School of Medicine, University of Southern California, Los Angeles, California, USA; Zilkha Neurogenetic Institute, Keck School of Medicine, University of Southern California, Los Angeles, California, USA; Department of Molecular Microbiology and Immunology, Keck School of Medicine, University of Southern California, Los Angeles, California, USA; Department of Translational Neuroscience, University Medical Center Utrecht, Utrecht University, Utrecht, The Netherlands

**Author notes:** Corresponding author. Please address correspondence to Prof. Justin Ichida.

**Keywords:** amyotrophic lateral sclerosis, frontotemporal dementia, apilimod, C9ORF72, NMDA-induced injury, endosomal trafficking, dipeptide repeat proteins, lysosomes, endosomes, glutamate receptors

## Abstract

The most common known cause of amyotrophic lateral sclerosis (ALS) and frontotemporal dementia (FTD) is a hexanucleotide repeat expansion (HRE) in *C9ORF72* that contributes to neurodegeneration by both loss-of-function (decreased C9ORF72 protein levels) and gain-of-function (e.g. dipeptide repeat protein production) mechanisms. Although therapeutics targeting the gain-of-function mechanisms are in clinical development, it is unclear if these will be efficacious given the contribution of C9ORF72 loss-of-function processes to neurodegeneration. Moreover, there is a lack of therapeutic strategies for *C9ORF72* ALS/FTD with demonstrated efficacy *in vivo*. Here, we show that small molecule inhibition of PIKFYVE kinase rescues both loss- and gain-of-function C9ORF72 disease mechanisms *in vivo*. We find that the reduction of C9ORF72 in mouse motor neurons leads to a decrease in early endosomes. In contrast, treatment with the PIKFYVE inhibitor apilimod increases the number of endosomes and lysosomes. We show that reduced C9ORF72 levels increases glutamate receptor levels in hippocampal neurons in mice, and that apilimod treatment rescues this excitotoxicity-related phenotype *in vivo*. Finally, we show that apilimod also alleviates the gain-of-function pathology induced by the *C9ORF72* HRE by decreasing levels of dipeptide repeat proteins derived from both sense and antisense *C9ORF72* transcripts in hippocampal neurons *in vivo*. Our data demonstrate the neuroprotective effect of PIKFYVE kinase inhibition in both gain- and loss-of-function murine models of *C9ORF72* ALS/FTD.

## Main text

The *C9ORF72* repeat expansion causes neurodegeneration through loss- and gain-of-function mechanisms [1–3]. Reduced C9ORF72 levels alter endosomal trafficking [3, 4], autophagy [5, 6], and lysosomes *in vitro* [3, 7–9], and increase glutamate receptors on neurons [3]. *C9ORF72* ALS/FTD (C9-ALS/FTD) gain-of-function mechanisms include neurotoxic dipeptide repeat proteins (DPRs) generated by repeat-associated non-AUG translation [3, 10–15].

Although antisense oligonucleotide therapeutics targeting sense *C9ORF72* transcripts are undergoing clinical testing, their efficacy may be limited given the contribution of C9ORF72 loss-of-function processes to neurodegeneration [3, 5, 16] and the fact that both sense and antisense *C9ORF72* transcripts generate DPRs. Problematically, there is a lack of other targets whose perturbation can rescue C9-ALS/FTD disease processes *in vivo*. We recently performed a phenotypic screen to identify small molecules that rescue the survival of C9-ALS/FTD induced motor neurons (iMNs) [3]. PIKFYVE kinase inhibitors such as apilimod potently rescued C9-ALS/FTD iMN survival [3]. Suppression of PIKFYVE using antisense oligonucleotides rescued C9-ALS iMN survival, confirming PIKFYVE as the therapeutic target [3].

PIKFYVE is a Class-III Phosphatidylinositol-5-kinase (PI5K) that synthesizes PI(3,5)P_2_ from PI3P [17]. PIKFYVE inhibition promotes endosomal maturation by increasing PI3P levels, and PI3P is also critical for autophagosome formation and engulfment of proteins designated for degradation [18, 19]. Therefore, PIKFYVE regulates cellular processes that are disrupted in C9-ALS/FTD, suggesting that altering PIKFYVE activity could modulate C9-ALS/FTD processes in patients. Here, we examine (1) the effects of the *C9ORF72* HRE and (2) the ability of PIKFYVE inhibition to rescue gain- and loss-of-function of *C9ORF72* ALS/FTD disease processes *in vivo*.

To determine if C9ORF72 deficiency alters endosomal trafficking *in vivo*, we used mice with approximately 50% reduced C9ORF72 levels in heterozygotes and a complete loss of C9ORF72 in homozygotes [20, 21]. We and others have shown that even modest changes in the number of intracellular vesicles such as endosomes, lysosomes, or autophagosomes by less than 50% can impair protein trafficking and degradation in neurons and reduce neuronal survival in neurodegenerative disease models [3, 22–25]. Moreover, mutations in one copy of *GBA*, which encodes a lysosomal enzyme, increase the risk of developing Parkinson’s disease by 20-30-fold [26]. Therefore, to increase the sensitivity of our analyses, we quantified endosome and lysosome numbers on a per cell basis in similar fashion to previous studies [3, 24, 25]. However, we included quantification of the median values and variation on a per animal basis to assess reliability of our results. We found that hippocampal neurons of *C9orf72^+/-^* mice, which mimic the ^~^50% reduction in C9ORF72 levels in C9-ALS/FTD patients, had fewer EEA1+ early endosomes than controls (Fig. 1A-C). Immunoblotting on total brain samples showed that PIKFYVE levels were similar between control and *C9orf72^-/-^* mice (Fig. 1D, E), indicating that differences in PIKFYVE levels did not cause the reduction in EEA1+ vesicles.

**Figure 1.**
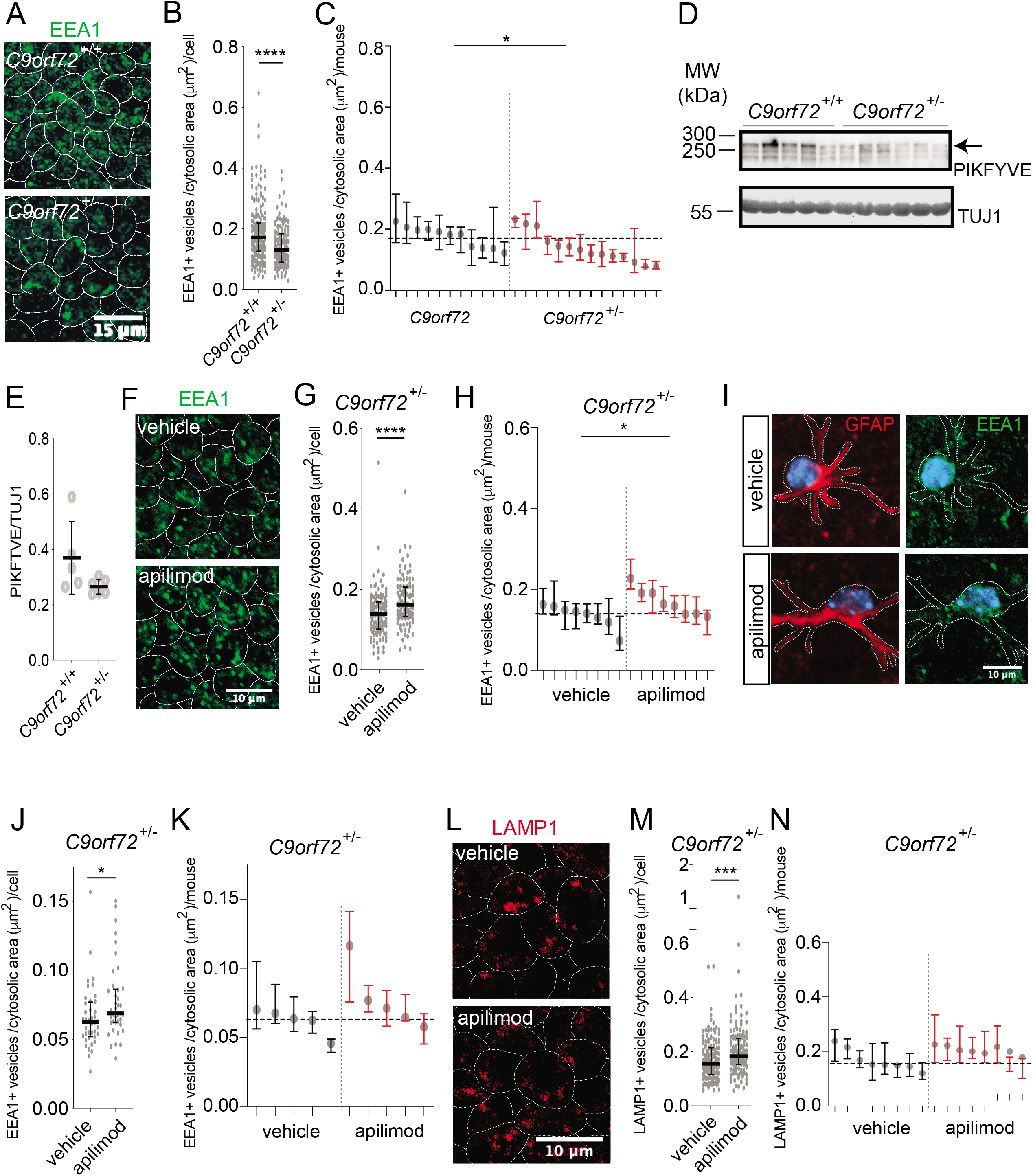
Endosome and lysosome changes induced by PIKFYVE inhibition in vivo oppose those caused by reduced C9orf72 levels. A. Images of EEA1+ vesicles in hippocampal neurons of C9orf72^+/+^ and C9orf72^+/-^ mice. Scale bar = 15 μm. B. EEA1+ vesicle number per μm^2^ of cytosolic space in TUJ1+ cells in C9orf72^+/+^ (n=195 cells from 11 mice) or C9orf72^+/-^ mice (n=132 cells from 14 mice). Median +/- interquartile range, Mann-Whitney test. C. EEA1+ vesicle number per μm^2^ of cytosolic space measured in TUJ1+ neurons of the hippocampus of C9orf72^+/+^ mice (n=11) and C9orf72^+/-^ mice (n=14). Median +/- interquartile range of cells for each mouse, unpaired t-test comparing mean values per mouse. Horizontal grey dotted line indicates the median of EEA1+ vesicle number per μm^2^ of cytosolic space measured in TUJ1+ neurons of the hippocampus of C9orf72^+/+^ mice. D. Western blot of PIKFYVE and TUJ1 levels in total brain samples from C9orf72^+/+^ and C9orf72^-/-^ mice. Each lane is a sample from a different mouse. E. Quantification of (D), PIKFYVE normalized to TUJ1 levels. Western blot samples from n=5 C9orf72^+/+^ mice and n=6 C9orf72^-/-^ mice. Mean +/- standard deviation, unpaired t-test. Outcome measure: PIKFYVE normalized to TUJ1 levels per mouse. F. Images of EEA1+ vesicle numbers in the hippocampus of C9orf72^+/-^ mice treated with apilimod or DMSO. Scale bar = 10 μm. G. EEA1+ vesicle number per μm^2^ of neuronal space measured in TUJ1+ cells in C9orf72^+/-^ mice treated by direct injection with apilimod (n=114 cells from 8 mice) or DMSO (vehicle control, n=118 cells from 8 mice) for 24 hours. Median +/- interquartile range, Mann-Whitney test. H. EEA1+ vesicle number per μm^2^ of neuronal space measured in TUJ1+ neurons of the hippocampus of C9orf72^+/-^ mice treated by direct injection with apilimod (n=8 mice) or DMSO (vehicle control, n=8 mice) for 24 hours per animal. Median +/- interquartile range of cells for each mouse, unpaired t-test comparing mean values per mouse. Horizontal grey dotted line indicates the median of EEA1+ vesicle number per μm^2^ of cytosolic space measured in TUJ1+ neurons of the hippocampus of vehicle-treated mice. I. Images of EEA1+ vesicle numbers in GFAP+ astrocytes in the hippocampus of C9orf72^+/-^ mice treated with apilimod or DMSO. Scale bar = 10 μm. J. EEA1+ vesicle number per μm^2^ of cytosolic space in GFAP+ astrocytes in the hippocampus of C9orf72^+/-^ mice treated by direct injection with apilimod (n=40 cells from 5 mice) or DMSO (vehicle control, n=40 cells from 5 mice) for 24 hours. Median +/- interquartile range, Mann-Whitney test. K. EEA1+ vesicle number per μm^2^ of cytosolic space measured in GFAP+ astrocytes of the dentate gyrus of C9orf72^+/-^ mice treated by direct injection with apilimod (n=5 mice) or DMSO (vehicle control, n=5 mice) for 24 hours per mouse. Median +/- interquartile range of cells for each mouse, unpaired t-test comparing mean values per mouse. Horizontal grey dotted line indicates the median of EEA1+ vesicle number per μm^2^ of cytosolic space measured in GFAP+ cells of the hippocampus of vehicle-treated mice. L. Images of LAMP1+ vesicle numbers in TUJ1+ cells in the hippocampus of C9orf72^+/-^ mice treated with apilimod or DMSO. Scale bar = 10 μm. M. LAMP1+ vesicle number per μm^2^ of cytosolic space in TUJ1+ cells in the hippocampus of C9orf72^-/+^ mice treated by direct injection with apilimod (n=132 cells from 8 mice) or DMSO (vehicle control, n=139 cells from 8 mice) for 24 hours. Median +/- interquartile range, Mann-Whitney test. N. LAMP1+ vesicle number per μm^2^ of cytosolic space measured in TUJ1+ neurons of the dentate gyrus of C9orf72^+/-^ mice treated by direct injection with apilimod (n=8 mice) or DMSO (vehicle control, n=8 mice) for 24 hours per mouse. Median +/- interquartile range of cells for each mouse, unpaired t-test comparing mean values per mouse. Horizontal grey dotted line indicates the median of LAMP1+ vesicle number per μm^2^ of cytosolic space measured in TUJ1+ neurons of the hippocampus of vehicle-treated mice. Dotted lines delineate cells. *p<0.05, **p<0.01, ***p<0.001, ****p<0.0001

It remains unclear whether the endosome and lysosome changes caused by C9ORF72 deficiency drive neurodegeneration or are adaptive compensatory changes that prevent neurodegeneration [3]. Because PIKFYVE inhibition promotes C9-ALS/FTD iMN survival [3], determining its effects on endosomes and lysosomes could elucidate this issue. Injection of apilimod directly to the hippocampus of *C9orf72^+/-^* mice increased the number of EEA1+ vesicles in both neurons and GFAP+ astrocytes (Fig. 1F-K).

C9ORF72-deficient human and mouse motor neurons have fewer LAMP1+ vesicles (lysosomes) than controls [3]. Apilimod increased LAMP1+ vesicle number in hippocampal neurons in *C9orf72^/-^* mice 24 hours after treatment (Fig. 1L-N). Thus, in hippocampal neurons in adult mice, PIKFYVE inhibition and C9ORF72 deficiency have opposing effects on endosomal and lysosomal numbers, suggesting that having fewer endosomes and/or lysosomes due to reduced C9ORF72 levels may drive neurodegeneration and increasing endosomes and/or lysosomes by PIKFYVE inhibition may rescue neurodegeneration.

C9ORF72-deficient mouse and human spinal motor neurons have increased glutamate receptor levels [3]. Because the *C9ORF72* HRE can induce neurodegeneration in the hippocampus, cortex, and spinal cord [27–30], we have used stereotactic injections into the hippocampus to measure the effects of C9ORF72 deficiency and PIKFYVE inhibition on NMDA-induced excitotoxicity [3]. Direct injection of NMDA into the hippocampus causes significantly greater excitotoxic injury in C9ORF72-deficient mice than controls [3].

To determine if the exacerbated excitotoxic injury in C9ORF72-deficient mice might result from increased glutamate receptor levels in brain regions affected by the *C9orf72* HRE, we measured NR1 (NMDA receptor subunit) and GLUR6/7 (kainate receptor subunit) levels in the hippocampus and frontal cortex of *C9orf72^+/+^* and *C9orf72^+/-^* mice. Changes in glutamate receptor levels by less than 50% [3, 31–33] and as little as 5% [34] can have a strong effect on neuronal function, sensitivity to neurotransmitters, and animal survival. Therefore, to increase the sensitivity of our analyses, we quantified glutamate receptor levels on a per cell basis in similar fashion to previous studies [3, 32, 33]. However, we included quantification of the median values and variation on a per animal basis to assess reliability of our results.

Immunohistochemistry showed that NR1 and GLUR6/7 levels were increased in the hippocampus and frontal cortex of *C9orf72^+/-^* mice compared to *C9orf72^+/+^* mice (Fig. 2A-F, J-O). To confirm that reduced C9ORF72 levels lead to increased glutamate receptor levels *in vivo*, we harvested post-synaptic densities from the brains of *C9orf72*^-/-^ mice, which have undetectable C9ORF72 levels [21]. Post-synaptic density preparations were enriched for PSD-95 and not enriched for presynaptic proteins including Synaptophysin and intracellular proteins including p53 (Fig. 2G). Immunoblotting confirmed that NR1 levels were increased in post-synaptic densities of *C9orf72^-/-^* mice, indicating they may affect neurotransmission (Fig. 2H, I). Thus, reduced C9ORF72 levels lead to increased glutamate receptor levels in multiple *C9ORF72* HRE-affected brain regions *in vivo*.

**Figure 2.**
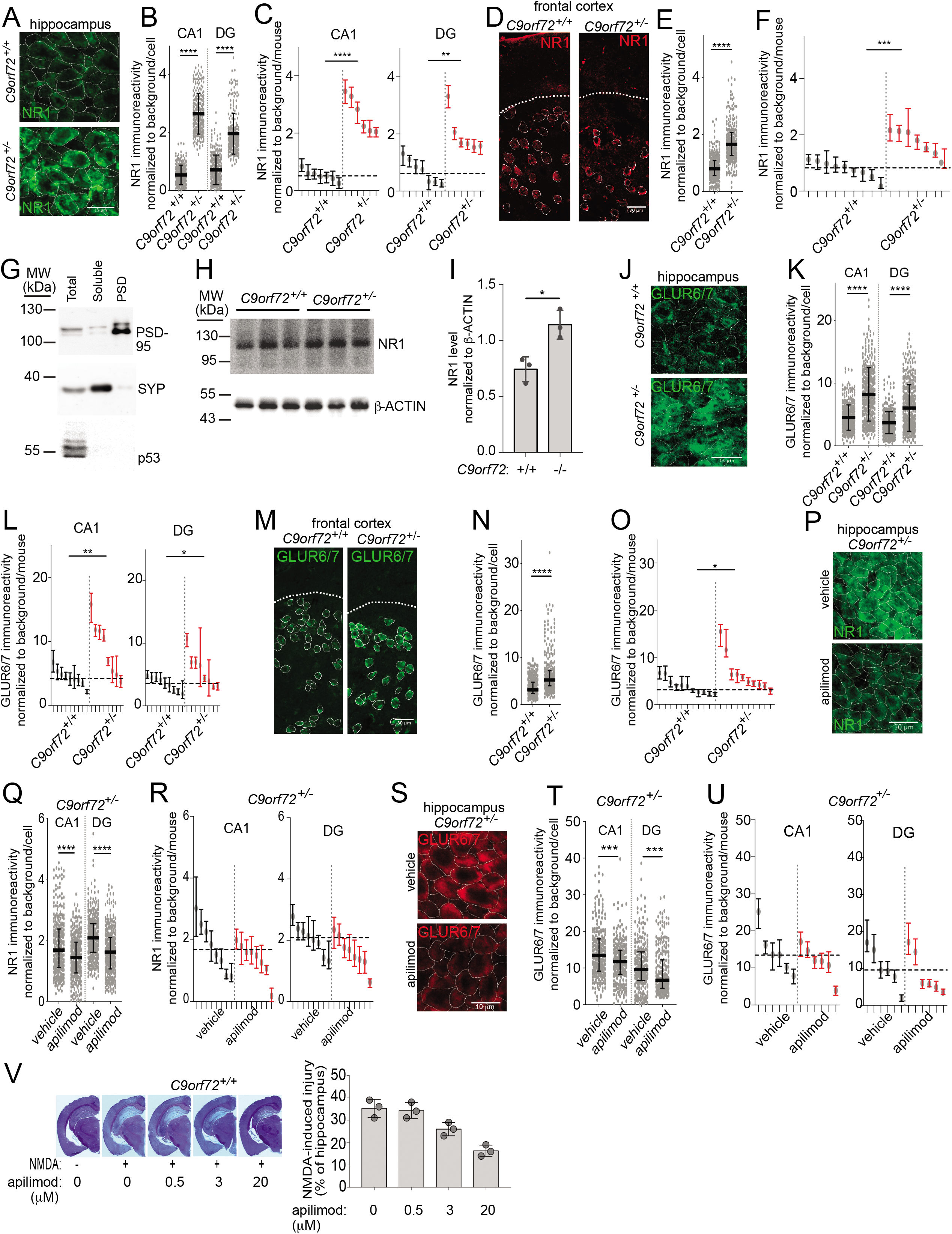
C9orf72 insufficiency increases glutamate receptor levels, which are rescued by apilimod. A. Images of NR1 levels in the hippocampus of C9orf72^+/+^ and C9orf72^+/-^ mice. Scale bar = 15 μm. B. Relative intensity of NR1 immunostaining in the hippocampus of C9orf72^+/+^ (n=280 cells per region from 7 mice) or C9orf72^+/-^ (n=240 cells per region from 6 mice) mice. Median +/- interquartile range, Mann-Whitney test comparing genotypes within each hippocampal region. DG = dentate gyrus C. Relative intensity of NR1 immunostaining in the hippocampus of C9orf72^+/+^ (n= 7 mice) or C9orf72^+/-^ (n= 6 mice) mice. Median +/- interquartile range of cells for each mouse, unpaired t-test comparing mean values per mouse. Horizontal grey dotted lines indicate the median of the relative intensity of NR1 immunostaining in the hippocampus of C9orf72^+/+^ mice. D. Images of NR1 levels in the frontal cortex of C9orf72^+/+^ and C9orf72^+/-^ mice. Scale bar = 30 μm. E. Relative intensity of NR1 immunostaining in the frontal cortex of C9orf72^+/+^ (n=321 cells from 9 mice) or C9orf72^+/-^ (n=262 cells from 7 mice) mice. Median +/- interquartile range, Mann-Whitney test. F. Relative intensity of NR1 immunostaining in frontal cortex of C9orf72^+/+^ (n= 9 mice) or C9orf72^+/-^ (n= 7 mice) mice. Median +/- interquartile range of cells for each mouse, unpaired t-test comparing mean values per mouse. Horizontal grey dotted line indicates the median of the relative intensity of NR1 immunostaining in the hippocampus of C9orf72^+/+^ mice. G. Western blot showing PSD-95, SYP, and p53 levels in the total, soluble (non-post-synaptic density), and post-synaptic density fractions derived from mouse brain. Levels of PSD-95 (post-synaptic density), Synaptophysin (SYP)(pre-synaptic), and p53 (intracellular) are examined in each fraction to measure the purity of the post-synaptic density fraction. H. Western blot showing NR1 and β-ACTIN levels in the post-synaptic density fractions of brain samples from C9orf72^+/+^ and C9orf72^-/-^ mice. Samples from n=3 mice per genotype. I. Quantification of (F), relative NR1 levels (normalized to β-ACTIN) in post-synaptic densities of brains of C9orf72^+/+^ or C9orf72^-/-^ mice, n=3 mice per genotype. Mean +/- s.e.m., unpaired t-test. J. Images of GLUR6/7 levels in the hippocampus of C9orf72^+/+^ and C9orf72^+/-^ mice. Scale bar = 15 μm. K. Relative intensity of GLUR6/7 immunostaining in the hippocampus of C9orf72^+/+^ (n=400 cells per region from 9 mice) or C9orf72^+/-^ (n=400 cells per region from 8 mice) mice. Median +/- interquartile range, Mann-Whitney test comparing genotypes within each hippocampal region. L. Relative intensity of GLUR6/7 immunostaining in the hippocampus of C9orf72^+/+^ (n= 9 mice) or C9orf72^+/-^ (n= 8 mice) mice. Median +/- interquartile range of cells for each mouse, unpaired t-test comparing mean values per mouse. Horizontal grey dotted lines indicate the median of the relative intensity of GLUR6/7 immunostaining in the hippocampus of C9orf72^+/+^ mice. M. Images of GLUR6/7 levels in the frontal cortex of C9orf72^+/+^ and C9orf72^+/-^ mice. Scale bar = 30 μm. N. Relative intensity of GLUR6/7 immunostaining in the frontal cortex C9orf72^+/+^ (n=331 cells from 11 mice) or C9orf72^+/-^ (n=306 cells from 10 mice) mice. Median +/- interquartile range, Mann-Whitney test. O. Relative intensity of GLUR6/7 immunostaining in the frontal cortex of C9orf72^+/+^ (n= 11 mice) or C9orf72^+/-^ (n= 10 mice) mice. Median +/- interquartile range of cells for each mouse, unpaired t-test comparing mean values per mouse. Horizontal grey dotted line indicates the median of the relative intensity of GLUR6/7 immunostaining in the frontal cortex of C9orf72^+/+^ mice. P. Images of NR1 levels in the hippocampus of C9orf72^+/-^ mice treated with apilimod or DMSO. Scale bar = 10 μm. Q. Relative intensity of NR1 immunostaining in the hippocampus of C9orf72^+/-^ mice treated by direct injection with apilimod (n=320 cells per region from 8 mice) or DMSO (vehicle control, n=320 cells per region from 8 mice) for 24 hours. Median +/- interquartile range, Mann-Whitney test comparing genotypes within the CA1 region. Mean +/- standard deviation, Unpaired t-test for DG region. R. Relative intensity of NR1 immunostaining in the hippocampus of C9orf72^+/-^ mice treated by direct injection with apilimod (n=8 mice) or DMSO (vehicle control, n=8 mice) for 24 hours. Median +/- interquartile range of cells for each mouse, unpaired t-test comparing mean values per mouse. Horizontal grey dotted lines indicate the median of the relative intensity of NR1 immunostaining in the hippocampus of vehicle-treated mice. S. Images of GLUR6/7 levels in the hippocampus of C9orf72^+/-^ mice treated with apilimod or DMSO. Scale bar = 10 μm. T. Relative intensity of GLUR6/7 immunostaining in the hippocampus of C9orf72^+/-^ mice treated by direct injection with apilimod (n=240 cells per region from 6 mice) or DMSO (vehicle control, n=240 cells per region from 6 mice) for 24 hours. Median +/- interquartile range, Mann-Whitney test. U. Relative intensity of GLUR6/7 immunostaining in the hippocampus of C9orf72^+/-^ mice treated by direct injection with apilimod (n=6 mice) or DMSO (vehicle control, n= 6 mice) for 24 hours. Median +/- interquartile range of cells for each mouse, unpaired t-test comparing mean values per mouse. Horizontal grey dotted lines indicate the median of the relative intensity of GLUR6/7 immunostaining in the hippocampus of vehicle-treated mice. V. Average hippocampal injury size in C9orf72^+/+^ mice when NMDA is co-injected with different doses of apilimod and incubated for 48 hours. n=3 brains per condition. Mean +/- std. dev., oneway ANOVA. White dotted lines indicate the extent of NMDA-induced hippocampal injury. Dotted lines delineate cells. *p<0.05, **p<0.01, ***p<0.001, ****p<0.0001

Apilimod rescues the increased susceptibility to NMDA-induced excitotoxicity in the hippocampus of *C9orf72^+/-^* mice [3]. However, it is unclear if this is accomplished in part through direct reversal of C9ORF72 loss-of-function disease processes, for example by lowering glutamate receptor levels, or through some unrelated mechanism that prevents cell death. In *C9orf72^+/-^* mice, apilimod treatment lowered NR1 and GLUR6/7 levels in the CA1 region and dentate gyrus (Fig. 2P-U). Thus, PIKFYVE inhibition can lower glutamate receptor levels in C9ORF72-deficient tissue. Consistent with the notion that apilimod-modulated NR1 levels would affect the NMDA-sensitivity of hippocampal neurons, apilimod dose-dependently reduced NMDA-induced neurodegeneration in the hippocampus after 48 hours (Fig. 2V). Thus, apilimod can rescue increased glutamate receptor levels, a C9ORF72 loss-of-function disease phenotype that increases susceptibility to glutamate-induced excitotoxicity.

C9-BAC mice harboring a human *C9ORF72* gene containing 100-1000 GGGGCC repeats recapitulate C9-ALS/FTD gain-of-function processes by producing DPRs that aggregate in neurons [35]. We previously showed that a single injection of apilimod decreased the number of poly-glycine-arginine (poly(GR))+ punctae in the dentate gyrus of C9-BAC mice [3]. However, poly(GR)+ DPRs are generated from the sense *C9ORF72* transcript, and we did not determine if PIKFYVE inhibition can reduce aggregates containing poly-proline-arginine (poly(PR)), which is made from the antisense *C9ORF72* transcript [36]. Additionally, we did not assess the effect of PIKFYVE inhibition on poly-glycine-proline (poly(GP))+ DPR aggregates, which are more prevalent than arginine-containing DPRs in C9-ALS/FTD patients and highly neurotoxic in mice [37]. Unlike the nuclear-localized poly(GR)+ and poly(PR)+ aggregates, poly(GP)+ aggregates are predominantly cytosolic [36], making it unclear if mechanisms that reduce poly(GR)+ aggregate levels would lower poly(GP)+ aggregates. Since multiple DPR species can cause neurotoxicity [38], a key measure of therapeutic efficacy against C9-ALS/FTD gain-of-function processes is the ability to diminish multiple DPRs.

Changes in DPR levels by less than 50% can have a strong effect on neuronal survival [3] and animal behavior [28]. Therefore, to increase the sensitivity of our analyses, we quantified DPR punctae levels on a per cell basis in similar fashion to previous studies [3, 15]. However, we included quantification of the median values and variation on a per animal basis to assess reliability of our results.

Apilimod increased the number of EEA1+ vesicles in C9-BAC mice after 48 hours (Fig. 3A-C), indicating that PIKFYVE inhibition results in similar endosomal changes in the presence or absence of the *C9ORF72* repeat expansion. 48 hours after injection into the hippocampus, apilimod reduced the number of poly(GP)+ (Fig. 3D-F) and poly(PR)+ (Fig. 3G-I) punctae in the dentate gyrus. Thus, increases in endosome numbers correlate with reduced DPR levels, and PIKFYVE inhibition can reduce nuclear and cytoplasmic DPR aggregates derived from both sense and antisense *C9ORF72* transcripts *in vivo*.

**Figure 3.**
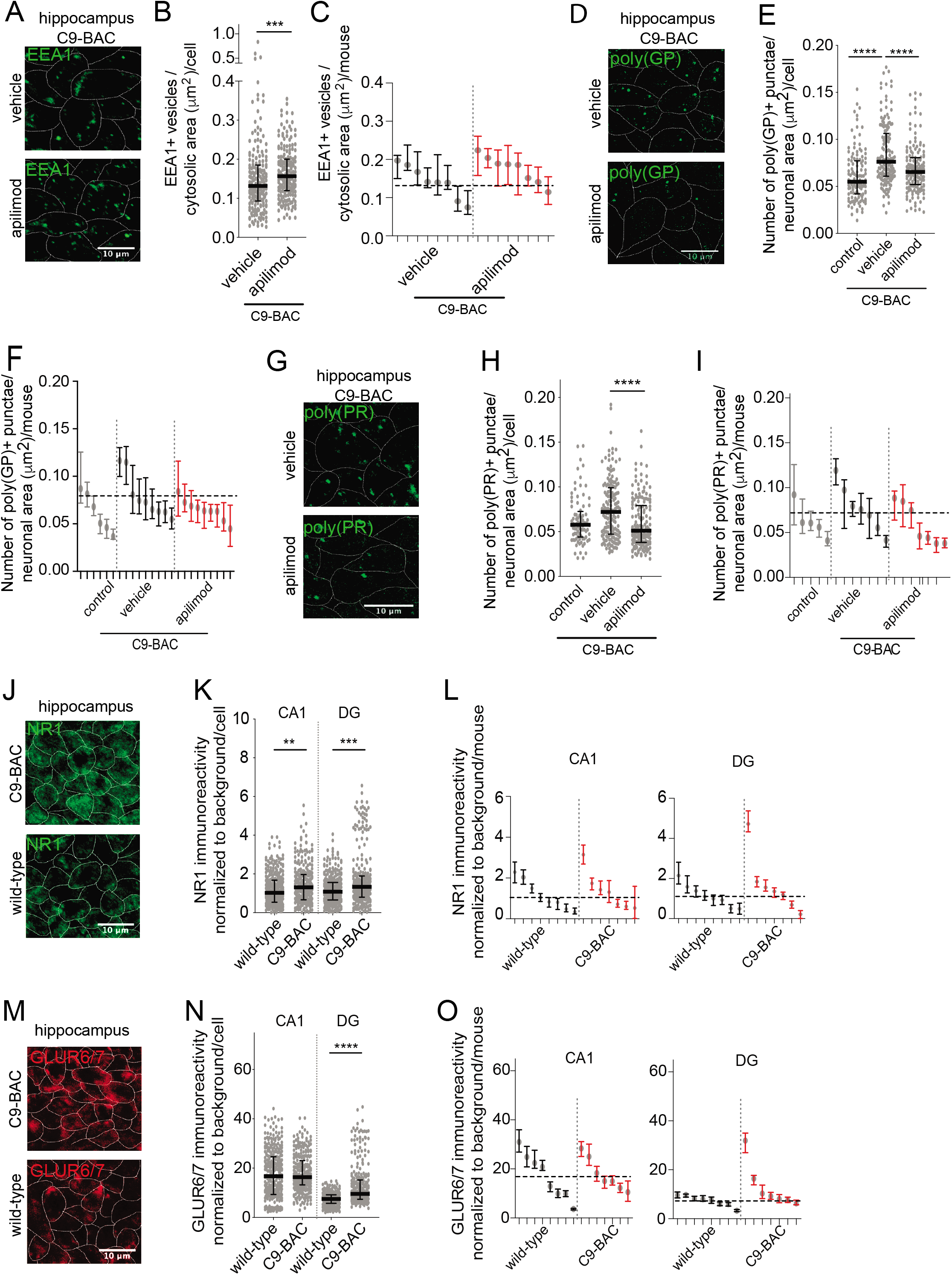
PIKFYVE inhibition rescues C9orf72 repeat expansion gain of function disease processes in vivo. A. Images of EEA1+ vesicles in the hippocampus of C9-BAC mice treated by direct injection with DMSO or apilimod. Scale bar = 10 μm. B. Number of EEA1+ vesicles per cytosolic area in the hippocampus of C9-BAC mice treated by direct injection with apilimod (n=186 cells from 8 mice) or DMSO (vehicle, n=185 cells from 8 mice) for 48 hours. Mean +/- standard deviation, unpaired t-test. C. Number of EEA1+ vesicles per cytosolic area in the hippocampus of C9-BAC mice treated by direct injection with apilimod (n=8 mice) or DMSO (vehicle, n= 8 mice) for 48 hours. Median +/- interquartile range of cells for each mouse, unpaired t-test comparing mean values per mouse. Horizontal grey dotted line indicates the median number of EEA1+ vesicles per cytosolic area in the hippocampus of C9-BAC mice treated by direct injection with DMSO. D. Images of poly(GP)+ punctae in the hippocampus of C9-BAC mice 48 hours after being treated by direct injection with DMSO or apilimod. Scale bar = 10 μm. E. Number of poly(GP)+ punctae per neuronal area in the hippocampus of C9-BAC mice treated by direct injection with apilimod (n=150 cells from 9 mice) or DMSO (vehicle, n=150 cells from 9 mice) for 48 hours. Control group, included for reference, are untreated wild-type mice (n=125 cells from 6 mice). Median +/- interquartile range, Mann-Whitney tests. F. Number of poly(GP)+ punctae per neuronal area in the hippocampus of C9-BAC mice treated by direct injection with apilimod (n=9 mice) or DMSO (vehicle, n=9 mice) for 48 hours. Control group, included for reference, are untreated wild-type mice (n=6 mice). Median +/- interquartile range of cells for each mouse, unpaired t-test comparing mean values per mouse between vehicle- and apilimod-treated conditions. Horizontal grey dotted line indicates the median number of poly(GP)+ punctae per neuronal area in the hippocampus of C9-BAC mice treated by direct injection with DMSO. G. Images of poly(PR)+ punctae in the hippocampus of C9-BAC mice 48 hours after being treated by direct injection with DMSO or apilimod. Scale bar = 10 μm. H. Number of poly(PR)+ punctae per neuronal area in the hippocampus of C9-BAC mice treated by direct injection with apilimod (n=160 cells from 7 mice) or DMSO (vehicle, n=160 cells from 7 mice) for 48 hours. Control group, included for reference, are untreated wild-type mice (n=84 cells from 5 mice). Median +/- interquartile range, Mann-Whitney test. I. Number of poly(PR)+ punctae per neuronal area in the hippocampus of C9-BAC mice treated by direct injection with apilimod (n=7 mice) or DMSO (vehicle, n=7 mice) for 48 hours. Control group, included for reference, are untreated wild-type mice (n=5 mice). Median +/- interquartile range of cells for each mouse, unpaired t-test comparing mean values per mouse between vehicle- and apilimod-treated conditions. Horizontal grey dotted line indicates the median number of poly(PR)+ punctae per neuronal area in the hippocampus of C9-BAC mice treated by direct injection with DMSO. J. Images of NR1 levels in the hippocampus of wild-type and C9-BAC mice. Scale bar = 10 μm. K. Relative intensity of NR1 immunostaining in wild-type and C9-BAC mice in hippocampal neurons of the CA1 or DG (n=320 cells per region from 8 mice (wild-type) and n=280 cells per region from 7 mice (C9-BAC)). Median +/- interquartile range, Mann-Whitney testing per region. L. Relative intensity of NR1 immunostaining in wild-type and C9-BAC mice in hippocampal neurons of the CA1 or DG (n=8 mice (wild-type) and n=7 mice (C9-BAC)). Median +/- interquartile range of cells for each mouse, unpaired t-test comparing mean values per mouse between wild-type and C9-BAC groups. Horizontal grey dotted lines indicate the median relative intensity of NR1 immunostaining in the hippocampus of wild-type mice. M. Images of GLUR6/7 levels in the hippocampus of wild-type and C9-BAC mice. Scale bar = 10 μm. N. Relative intensity of GLUR6/7 immunostaining in wild-type and C9-BAC mice in hippocampal neurons of the CA1 or DG (n=320 cells per region from 8 mice (wild-type) and n=280 cells per region from 7 mice (C9-BAC)). Median +/- interquartile range, Mann-Whitney testing per region. O. Relative intensity of GLUR6/7 immunostaining in wild-type and C9-BAC mice in hippocampal neurons of the CA1 or DG (n=8 mice (wild-type) and n=7 mice (C9-BAC)). Median +/- interquartile range of cells for each mouse, unpaired t-test comparing mean values per mouse between wild-type and C9-BAC groups. Horizontal grey dotted lines indicate the median relative intensity of GLUR6/7 immunostaining in the hippocampus of wild-type mice. Dotted lines delineate cells. *p<0.05, **p<0.01, ***p<0.001, ****p<0.0001

Surprisingly, C9-BAC mice contained slightly elevated hippocampal NR1 and GLUR6/7 levels (Fig. 3J-O), indicating that gain- and loss-of-function *C9ORF72* processes can both lead to increased glutamate receptor levels. Thus, C9-ALS/FTD gain- and loss-of-function processes can have similar effects on glutamate receptor levels *in vivo*, reinforcing the importance of curtailing both gain- and loss-of-function processes.

Here, we show that C9ORF72 insufficiency leads to fewer endosomes and increased surface-bound glutamate receptor levels in mice, which could sensitize neurons to glutamate-induced excitotoxicity. PIKFYVE inhibition reverses endosome, lysosome, and glutamate receptor changes induced by reduced C9ORF72 function in mice. It also significantly reduces both cytosolic and nuclear, and both sense and antisense transcript-derived DPR aggregates in hippocampal neurons. Thus, our data provide some of the first evidence that a pharmacological intervention, PIKFYVE inhibition, can rescue both gain- and loss-of-function C9ORF72 disease processes *in vivo*.

## Methods

### Animal Care

All animal use and care were in accordance with local institution guidelines of the University Medical Center Utrecht and approved by the Dierexperimenten Ethische Commissie Utrecht with the protocol number DEC 2013.I.09.069. Wild-type C57BL6/J (strain: 000664), C9ORF72 KO (C57BL/6J-3110043021Rikem5Lutzy/J, strain: 027068), and C9-BAC (C57BL/6J-Tg(C9orf72_i3)112Lutzy/J, strain: 023099) were purchased from Jackson Laboratories. Mice were housed in standard conditions with food and water ad libitum in the conventional vivarium at the University of Southern California. All animal use and care were in accordance with local institution guidelines of the University of Southern California and the IACUC board of the University of Southern California with the protocol numbers 20546 and 11938.

### Direct injection of apilimod

Mice were anesthetized with i.p. ketamine (100 mg/kg) and xylazine (10 mg/kg), and body temperature kept at 36.9 ± 0.1 °C with a thermostatic heating pad. Mice were placed in a stereotactic apparatus (ASI Instruments, USA) and the head is fixed accordingly. A burr hole was drilled, and an injection needle (33 gauge) was lowered into the hippocampus between CA1 and the dentate gyrus (AP −2.0, ML +1.5, DV −1.8). Apilimod (0.3 μl of 0.5, 3, or 20 μM in phosphate-buffered saline, pH 7.4) was infused over 2 min using a micro-injection system (World Precision Instruments). The needle was left in place for an additional 8 min after the injection. Animals were euthanized 24h-48h later (as stated in the text).

### Immunohistochemistry

*C9orf72*^+/-^, C9-BAC, and wildtype controls were transcardially perfused with phosphate buffered saline (PBS) and subsequently with 4% formaldehyde. Cryoprotection occurred in 20% sucrose. After snap freezing, tissue was sectioned by cryostat at 20 μm thickness and stained with the following primary antibodies: NR1 (MAB363, EMD Millipore), GLUR6/7 (04-921, EMD Millipore), EEA1 (sc-6415, N19, G4, Santa Cruz), LAMP1 (ab24170, Abcam), NeuN (MAB374, EMD Millipore), ChAT (AB144P, Millipore Sigma), poly(PR) (23979-1-AP, ProteinTech), and poly(GP) (24494-1-AP, ProteinTech). Antigen retrieval by sodium citrate occurred before NR1 staining, and with Target Retrieval Solution (pH 9, Agilent, Dako) for the poly(GP) staining. Images were collected using a Zeiss LSM780 or LSM800 confocal microscope. Glutamate subunit intensity measurements occurred with ImageJ where the mean cytosolic intensity was subtracted by a background measurement collected near to the measured neuron, and then divided by the background measurement. The scientist performing the glutamate receptor subunit intensity, EEA1 vesicle, LAMP1 vesicle, poly(PR)+ punctae, and poly(GP)+ punctae quantification was blinded to the genotype or treatment condition of the samples.

Sections used for staining and quantification from all experiments were a random selection from across the hippocampus. For images of EEA1+ and LAMP1+ vesicles the images were adjusted for publication to ensure that the background intensity was similar. Confocal microscopy images of poly(GP)+ or poly(PR)+ punctae were adjusted for brightness and contrast for optimal visualization of the poly(GP)+ or poly(PR)+ punctae; this occurred to the same extent for the vehicle- and apilimod-treated side of the hippocampus. No corrections were applied to the images of glutamate receptor stainings, as these are based on intensity of signal measurements.

### Western Blot

Mouse spinal cord tissue was collected in RIPA buffer (Sigma-Aldrich) with a protease inhibitor cocktail (Roche). Membrane samples were obtained by the Plasma Membrane Isolation Kit (Abcam) according to the manufacturer’s instructions. Postsynaptic density extraction occurred by a protocol published previously [39]. In brief, tissues were homogenized in cold Sucrose Buffer (320 mM Sucrose, 10 mM HEPES pH 7.4, 2 mM EDTA, 30 mM NaF, 40 mM β-Glycerophosphate, 10 mM Na3VO4, and protease inhibitor cocktail (Roche)) using a tissue grinder and then spun down at 500 g for 6 min at 4 °C. The supernatant was re-centrifuged at 10,000 g for 10 min at 4 °C. The supernatant was collected as the ‘Total’ fraction, and the pellet was resuspended in cold Triton buffer (50 mM HEPES pH 7.4, 2 mM EDTA, 50 mM NaF, 40 mM β-Glycerophosphate, 10 mM Na3VO4, 1% Triton X-100 and protease inhibitor cocktail (Roche)) and then spun down at 30,000 RPM using a Beckman rotor MLA-130 for 40 min at 4 °C. The supernantant was collected as the ‘Triton’ fraction and the pellet was resuspended in DOC buffer (50 mM HEPES pH 9.0, 50 mM NaF, 40 mM β-Glycerophosphate, 10 mM Na3VO4, 20 μM ZnCl2, 1% sodium deoxycholate and protease inhibitor cocktail (Roche)) and collected as the “DOC”, PSD-enriched fraction. Protein quantity was measured by the BCA assay (Pierce) and samples were run on a 10% SDS gel at 4 °C. The gel was transferred onto PVDF membrane (GE Healthcare) using Trans-Blot Semi-Dry transfer cell (Biorad). The membrane was blocked with 5% milk in 0.1% PBS-Tween 20 (PBS-T) (Sigma-Aldrich), incubated with primary antibodies overnight at 4 °C, washed three times with 0.1% PBS-T, then incubated with horseradish peroxidase (HRP)-conjugated secondary antibody (Santa Cruz). After three washes with 0.1% PBS-T, blots were visualized using an Amersham ECL Western Blotting Detection Kit (GE) or the SuperSignal West Femto Maximum Sensitivity Substrate (Thermo) and developed on X-ray film (Genesee Scientific). The following primary antibodies were used: goat anti-Actin-HRP (Santa Cruz, cat. no. sc-47778 HRP, 1:4,000), mouse anti-NR1 (Novus, cat. no. NB300118, 1:2,000), mouse anti-PSD-95 (Thermo, cat. no. MA1-045, 1:1,000), mouse anti-p53 (Cell Signaling, cat. no. 2524S, 1:1,000), mouse anti-Synaptophysin (Sigma, cat. no. S5768, 1:1000), anti-mouse HRP (Cell Signaling, cat. no. 7076S, 1:5,000), and anti-rabbit HRP (Cell Signaling, cat. no. 7074S, 1:5,000).

### Statistics

Analysis was performed with the statistical software package Prism Origin v.7.0a and v8.0a (GraphPad Software). The normal distribution of datasets was tested by the D’Agostino-Pearson omnibus normality test. Differences between two groups were analyzed using a two-tailed Student’s t test, unless the data was non-normally distributed for which two-sided Mann-Whitney testing was used. Differences between more than two groups were analyzed by one way-ANOVA with Tukey correction for multiple testing unless the data was non-normally distributed for which Kruskal-Wallis testing was used. Mean and standard deviation or standard error of the mean were used for normally distributed datasets, and the median and interquartile range were used for not normally distributed datasets. Significance was assumed at p < 0.05. Details of the statistical analyses for each panel are included in the figure legends.

## List of abbreviations

ALS: amyotrophic lateral sclerosis
CA: Cornu Ammonis
C9-ALS/FTD: C9orf72 ALS/FTD
C9-BAC mice: C9orf72 containing 100-1000 GGGGCC repeats mice
DG: dentate gyrus
DPR: dipeptide repeat protein
EEA1: Early Endosome Antigen 1
FTD: frontotemporal dementia
GP: glycine-proline repeat DPR
GR: glycine-arginine repeat DPR
HRE: hexanucleotide repeat expansion
HRP: horseradish peroxidase
iPSC: induced pluripotent stem cell
iMN: induced motor neuron
LAMP1: Lysosomal Associated Membrane Protein 1
NMDA: N-methyl-D-aspartate
PBS: phosphate buffered saline
PI5K: Phosphatidylinositol 5-kinase
PR: proline-arginine repeat DPR
RAN: repeat-associated non-AUG

## Declarations

### Ethics approval

All animal use and care were in accordance with local institution guidelines of the University of Southern California and the IACUC board of the University of Southern California with the protocol numbers 20546 and 11938.

### Consent for publication

Not applicable.

### Availability of data and material

The datasets used and/or analysed during the current study are available from the corresponding author on reasonable request.

### Authors’ contributions

KAS designed and planned the experiments, performed the initial experiments, supervised experiments, analyzed the data, and wrote the manuscript. YL identified the PIKFYVE target. CS, AS, NK, DK, KAG, and YL performed the experiments, and quantified readouts. YW performed the direct injection experiments, and MC performed drug treatment surgeries (unpublished results) that enabled the study. SL and WHC performed post-synaptic density isolations and performed all Western blot experiments. RJP, KS and VRV provided tissue, and supervised the work. PC and BZ designed experiments, and supervised the work. YS and YL provided unpublished results that enabled the study. JKI conceived the study, designed and supervised the experiments, and wrote the manuscript. All authors agreed with the final version of the manuscript.

## Acknowledgements

We thank the lab members of the Ichida lab for their useful feedback, and the University of Southern California Department of Animal Resources staff for their outstanding care.

## References

1. DeJesus-Hernandez M, Mackenzie IR, Boeve BF, Boxer AL, Baker M, Rutherford NJ, Nicholson AM, Finch NA, Flynn H, Adamson J et al: Expanded GGGGCC hexanucleotide repeat in noncoding region of C9ORF72 causes chromosome 9p-linked FTD and ALS. Neuron 2011, 72(2):245–256.

2. Renton AE, Majounie E, Waite A, Simon-Sanchez J, Rollinson S, Gibbs JR, Schymick JC, Laaksovirta H, van Swieten JC, Myllykangas L et al: A hexanucleotide repeat expansion in C9ORF72 is the cause of chromosome 9p21-linked ALS-FTD. Neuron 2011, 72(2):257–268.

3. Shi Y, Lin S, Staats KA, Li Y, Chang WH, Hung ST, Hendricks E, Linares GR, Wang Y, Son EY et al: Haploinsufficiency leads to neurodegeneration in C9ORF72 ALS/FTD human induced motor neurons. Nat Med 2018, 24(3):313–325.

4. Farg MA, Sundaramoorthy V, Sultana JM, Yang S, Atkinson RA, Levina V, Halloran MA, Gleeson PA, Blair IP, Soo KY et al: C9ORF72, implicated in amytrophic lateral sclerosis and frontotemporal dementia, regulates endosomal trafficking. Hum Mol Genet 2014, 23(13):3579–3595.

5. Sellier C, Campanari ML, Julie Corbier C, Gaucherot A, Kolb-Cheynel I, Oulad-Abdelghani M, Ruffenach F, Page A, Ciura S, Kabashi E et al: Loss of C9ORF72 impairs autophagy and synergizes with polyQ Ataxin-2 to induce motor neuron dysfunction and cell death. EMBO J 2016.

6. Sullivan PM, Zhou X, Robins AM, Paushter DH, Kim D, Smolka MB, Hu F: The ALS/FTLD associated protein C9orf72 associates with SMCR8 and WDR41 to regulate the autophagy-lysosome pathway. Acta neuropathologica communications 2016, 4(1):51.

7. Zhang Y, Burberry A, Wang JY, Sandoe J, Ghosh S, Udeshi ND, Svinkina T, Mordes DA, Mok J, Charlton M et al: The C9orf72-interacting protein Smcr8 is a negative regulator of autoimmunity and lysosomal exocytosis. Genes Dev 2018, 32(13-14):929–943.

8. Amick J, Roczniak-Ferguson A, Ferguson SM: C9orf72 binds SMCR8, localizes to lysosomes, and regulates mTORC1 signaling. Mol Biol Cell 2016, 27(20):3040–3051.

9. Corrionero A, Horvitz HR: A C9orf72 ALS/FTD Ortholog Acts in Endolysosomal Degradation and Lysosomal Homeostasis. Curr Biol 2018, 28(10):1522–1535 e1525.

10. Cooper-Knock J, Higginbottom A, Stopford MJ, Highley JR, Ince PG, Wharton SB, Pickering-Brown S, Kirby J, Hautbergue GM, Shaw PJ: Antisense RNA foci in the motor neurons of C9ORF72-ALS patients are associated with TDP-43 proteinopathy. Acta Neuropathol 2015, 130(1):63–75.

11. Donnelly CJ, Zhang PW, Pham JT, Haeusler AR, Mistry NA, Vidensky S, Daley EL, Poth EM, Hoover B, Fines DM et al: RNA toxicity from the ALS/FTD C9ORF72 expansion is mitigated by antisense intervention. Neuron 2013, 80(2):415–428.

12. Gendron TF, Bieniek KF, Zhang YJ, Jansen-West K, Ash PE, Caulfield T, Daughrity L, Dunmore JH, Castanedes-Casey M, Chew J et al: Antisense transcripts of the expanded C9ORF72 hexanucleotide repeat form nuclear RNA foci and undergo repeat-associated non-ATG translation in c9FTD/ALS. Acta Neuropathol 2013, 126(6):829–844.

13. Mori K, Weng SM, Arzberger T, May S, Rentzsch K, Kremmer E, Schmid B, Kretzschmar HA, Cruts M, Van Broeckhoven C et al: The C9orf72 GGGGCC repeat is translated into aggregating dipeptide-repeat proteins in FTLD/ALS. Science 2013, 339(6125):1335–1338.

14. Sareen D, O’Rourke JG, Meera P, Muhammad AK, Grant S, Simpkinson M, Bell S, Carmona S, Ornelas L, Sahabian A et al: Targeting RNA foci in iPSC-derived motor neurons from ALS patients with a C9ORF72 repeat expansion. Sci Transl Med 2013, 5(208):208ra149.

15. Wen X, Tan W, Westergard T, Krishnamurthy K, Markandaiah SS, Shi Y, Lin S, Shneider NA, Monaghan J, Pandey UB et al: Antisense Proline-Arginine RAN Dipeptides Linked to C9ORF72-ALS/FTD Form Toxic Nuclear Aggregates that Initiate In Vitro and In Vivo Neuronal Death. Neuron 2014, 84(6):1213–1225.

16. Ciura S, Lattante S, Le Ber I, Latouche M, Tostivint H, Brice A, Kabashi E: Loss of function of C9orf72 causes motor deficits in a zebrafish model of amyotrophic lateral sclerosis. Ann Neurol 2013, 74(2):180–187.

17. Lemmon MA: Membrane recognition by phospholipid-binding domains. Nat Rev Mol Cell Biol 2008, 9(2):99–111.

18. Martin S, Harper CB, May LM, Coulson EJ, Meunier FA, Osborne SL: Inhibition of PIKfyve by YM-201636 dysregulates autophagy and leads to apoptosis-independent neuronal cell death. PLoS One 2013, 8(3):e60152.

19. Seebohm G, Neumann S, Theiss C, Novkovic T, Hill EV, Tavare JM, Lang F, Hollmann M, Manahan-Vaughan D, Strutz-Seebohm N: Identification of a novel signaling pathway and its relevance for GluA1 recycling. PLoS One 2012, 7(3):e33889.

20. Koppers M, Blokhuis AM, Westeneng HJ, Terpstra ML, Zundel CA, Vieira de Sa R, Schellevis RD, Waite AJ, Blake DJ, Veldink JH et al: C9orf72 ablation in mice does not cause motor neuron degeneration or motor deficits. Ann Neurol 2015.

21. O’Rourke JG, Bogdanik L, Yanez A, Lall D, Wolf AJ, Muhammad AK, Ho R, Carmona S, Vit JP, Zarrow J et al: C9orf72 is required for proper macrophage and microglial function in mice. Science 2016, 351(6279):1324–1329.

22. Schwenk BM, Hartmann H, Serdaroglu A, Schludi MH, Hornburg D, Meissner F, Orozco D, Colombo A, Tahirovic S, Michaelsen M et al: TDP-43 loss of function inhibits endosomal trafficking and alters trophic signaling in neurons. EMBO J 2016, 35(21):2350–2370.

23. Dehay B, Bove J, Rodriguez-Muela N, Perier C, Recasens A, Boya P, Vila M: Pathogenic lysosomal depletion in Parkinson’s disease. J Neurosci 2010, 30(37):12535–12544.

24. Webster CP, Smith EF, Bauer CS, Moller A, Hautbergue GM, Ferraiuolo L, Myszczynska MA, Higginbottom A, Walsh MJ, Whitworth AJ et al: The C9orf72 protein interacts with Rab1a and the ULK1 complex to regulate initiation of autophagy. EMBO J 2016, 35(15):1656–1676.

25. Yang M, Liang C, Swaminathan K, Herrlinger S, Lai F, Shiekhattar R, Chen JF: A C9ORF72/SMCR8-containing complex regulates ULK1 and plays a dual role in autophagy. Science advances 2016, 2(9):e1601167.

26. Migdalska-Richards A, Schapira AH: The relationship between glucocerebrosidase mutations and Parkinson disease. J Neurochem 2016, 139 Suppl 1:77–90.

27. Pletnikova O, Sloane KL, Renton AE, Traynor BJ, Crain BJ, Reid T, Zu T, Ranum LP, Troncoso JC, Rabins PV et al: Hippocampal sclerosis dementia with the C9ORF72 hexanucleotide repeat expansion. Neurobiol Aging 2014, 35(10):2419 e2417–2421.

28. Jiang J, Zhu Q, Gendron TF, Saberi S, McAlonis-Downes M, Seelman A, Stauffer JE, Jafar-Nejad P, Drenner K, Schulte D et al: Gain of Toxicity from ALS/FTD-Linked Repeat Expansions in C9ORF72 Is Alleviated by Antisense Oligonucleotides Targeting GGGGCC-Containing RNAs. Neuron 2016, 90(3):535–550.

29. Simon-Sanchez J, Dopper EG, Cohn-Hokke PE, Hukema RK, Nicolaou N, Seelaar H, de Graaf JR, de Koning I, van Schoor NM, Deeg DJ et al: The clinical and pathological phenotype of C9ORF72 hexanucleotide repeat expansions. Brain 2012, 135(Pt 3):723–735.

30. Hsiung GY, DeJesus-Hernandez M, Feldman HH, Sengdy P, Bouchard-Kerr P, Dwosh E, Butler R, Leung B, Fok A, Rutherford NJ et al: Clinical and pathological features of familial frontotemporal dementia caused by C9ORF72 mutation on chromosome 9p. Brain 2012, 135(Pt 3):709–722.

31. Selvaraj BT, Livesey MR, Zhao C, Gregory JM, James OT, Cleary EM, Chouhan AK, Gane AB, Perkins EM, Dando O et al: C9ORF72 repeat expansion causes vulnerability of motor neurons to Ca(2+)-permeable AMPA receptor-mediated excitotoxicity. Nat Commun 2018, 9(1):347.

32. Chen P, Gu Z, Liu W, Yan Z: Glycogen synthase kinase 3 regulates N-methyl-D-aspartate receptor channel trafficking and function in cortical neurons. Mol Pharmacol 2007, 72(1):40–51.

33. Wei J, Liu W, Yan Z: Regulation of AMPA receptor trafficking and function by glycogen synthase kinase 3. J Biol Chem 2010, 285(34):26369–26376.

34. Mohn AR, Gainetdinov RR, Caron MG, Koller BH: Mice with reduced NMDA receptor expression display behaviors related to schizophrenia. Cell 1999, 98(4):427–436.

35. O’Rourke JG, Bogdanik L, Muhammad AK, Gendron TF, Kim KJ, Austin A, Cady J, Liu EY, Zarrow J, Grant S et al: C9orf72 BAC Transgenic Mice Display Typical Pathologic Features of ALS/FTD. Neuron 2015, 88(5):892–901.

36. Freibaum BD, Taylor JP: The Role of Dipeptide Repeats in C9ORF72-Related ALS-FTD. Frontiers in molecular neuroscience 2017, 10:35.

37. Chew J, Gendron TF, Prudencio M, Sasaguri H, Zhang YJ, Castanedes-Casey M, Lee CW, Jansen-West K, Kurti A, Murray ME et al: Neurodegeneration. C9ORF72 repeat expansions in mice cause TDP-43 pathology, neuronal loss, and behavioral deficits. Science 2015, 348(6239):1151–1154.

38. Lee YB, Baskaran P, Gomez-Deza J, Chen HJ, Nishimura AL, Smith BN, Troakes C, Adachi Y, Stepto A, Petrucelli L et al: C9orf72 poly GA RAN-translated protein plays a key role in amyotrophic lateral sclerosis via aggregation and toxicity. Hum Mol Genet 2017, 26(24):4765–4777.

39. Li J, Wilkinson B, Clementel VA, Hou J, O’Dell TJ, Coba MP: Long-term potentiation modulates synaptic phosphorylation networks and reshapes the structure of the postsynaptic interactome. Sci Signal 2016, 9(440):rs8.

